# Nucleoli as drivers of nuclear remodelling in cardiomyocytes during Heart Failure

**DOI:** 10.64898/2026.02.24.707499

**Authors:** Ingrid Matzer, Houcheng Wang, Aleksandra N. Kozyrina, Jiarong Fu, Thomas Iskratsch, Massimo Vassalli, Senka Ljubojevic-Holzer, Julia Gorelik, Pamela Swiatlowska

**Author notes:** First Author (Ingrid Matzer, Houcheng Wang).

## Abstract

Cardiomyocyte mechanotransduction has traditionally focused on the sarcomere and cytoskeleton, yet emerging evidence highlights the nucleus as an active mechanical responder. To adapt to the dynamic mechanical environment, the nucleus forms nuclear invaginations (NIs), double-membrane folds that provide structural support to chromatin and incorporate nuclear pore complexes to facilitate nucleo–cytoplasmic transport, including Ca^2+^ transport. However, how these structures are formed is not yet understood. We leveraged advances in high-resolution microscopy, mechanical stimulation, rat models and human samples, to study the formation, function and remodelling of cardiac NIs in Heart Failure (HF). Here, we demonstrate that the formation of NI in cardiomyocytes is regulated by both, cytoskeleton such as actin and detyrosinated microtubules as well as intranuclear nucleolar interactions with NI disruption resulting in elevated baseline nuclear Ca^2+^. In a 16-week post–Myocardial Infarction (MI) end-stage HF rat model, as well as in human Dilated Cardiomyopathy samples, a marked reduction in NIs was observed. Importantly, NI loss was already evident at 8 weeks post-MI, preceding detectable cytoskeletal stiffening. At this stage, we observed increased DNA damage in the peri-nucleolar region, accompanied by nucleolar remodelling and a shift in nucleolar biomechanical properties. In conclusion, preserving nucleolar integrity emerges as a potential target for intervention.

## Introduction

The sarcomere and the cytoskeleton have been identified as major cardiomyocyte actors in the mechanotransduction process, but a growing evidence is emerging that the nucleus can also respond to mechanical stimuli through structural adaptation and specialised structures such as the LINC complex (1).

One key nuclear adaptation is the formation of nuclear invaginations (NIs), deep double-membrane folds that extend into the nucleoplasm and function as nuclear T-tubules. Previous reports show that in Heart Failure (HF), nucleoplasmic Ca^2+^ changes precede cytoplasmic Ca^2+^ alterations and that NIs serve as entry routes for Ca^2+^ into the nucleus, underscoring the need to understand how NIs remodel under pathological conditions (2). However, the regulatory mechanisms of NIs, and how they are modified in HF with shifting mechanical conditions, remain largely unclear.

## Results and Conclusions

Using adult rat left ventricle cardiomyocytes and high-resolution imaging, we first dissected the regulation of NIs under physiological conditions. Pharmacological disruption of actin filaments with cytochalasin D (CytD) markedly affected NIs number, whereas microtubule disruption had no detectable impact. In contrast, treatment with parthenolide (PTL) to reduce detyrosinated microtubules (detyrMT), which are key determinants of cardiomyocyte mechanical properties and extend into the NIs, led to a decrease of NIs (Fig.Ai) (3). Using line-scan confocal microscopy combined with Fluo-4AM dye, we assessed nuclear Ca^2+^ (nCa^2+^) handling as a readout of nucleo– cytoplasmic communication. While cytoplasmic Ca^2+^ transients were unchanged after either treatment, baseline nCa^2+^ levels rose, particularly at higher pacing frequencies that mimic increased cardiac workload. Because NIs contain nuclear pores and reduce diffusion distances, their loss disrupts nCa^2+^ homeostasis. These findings establish that NIs are active organizers of nuclear signalling rather than passive features (Fig.Aii-iii).

**Figure 1.**
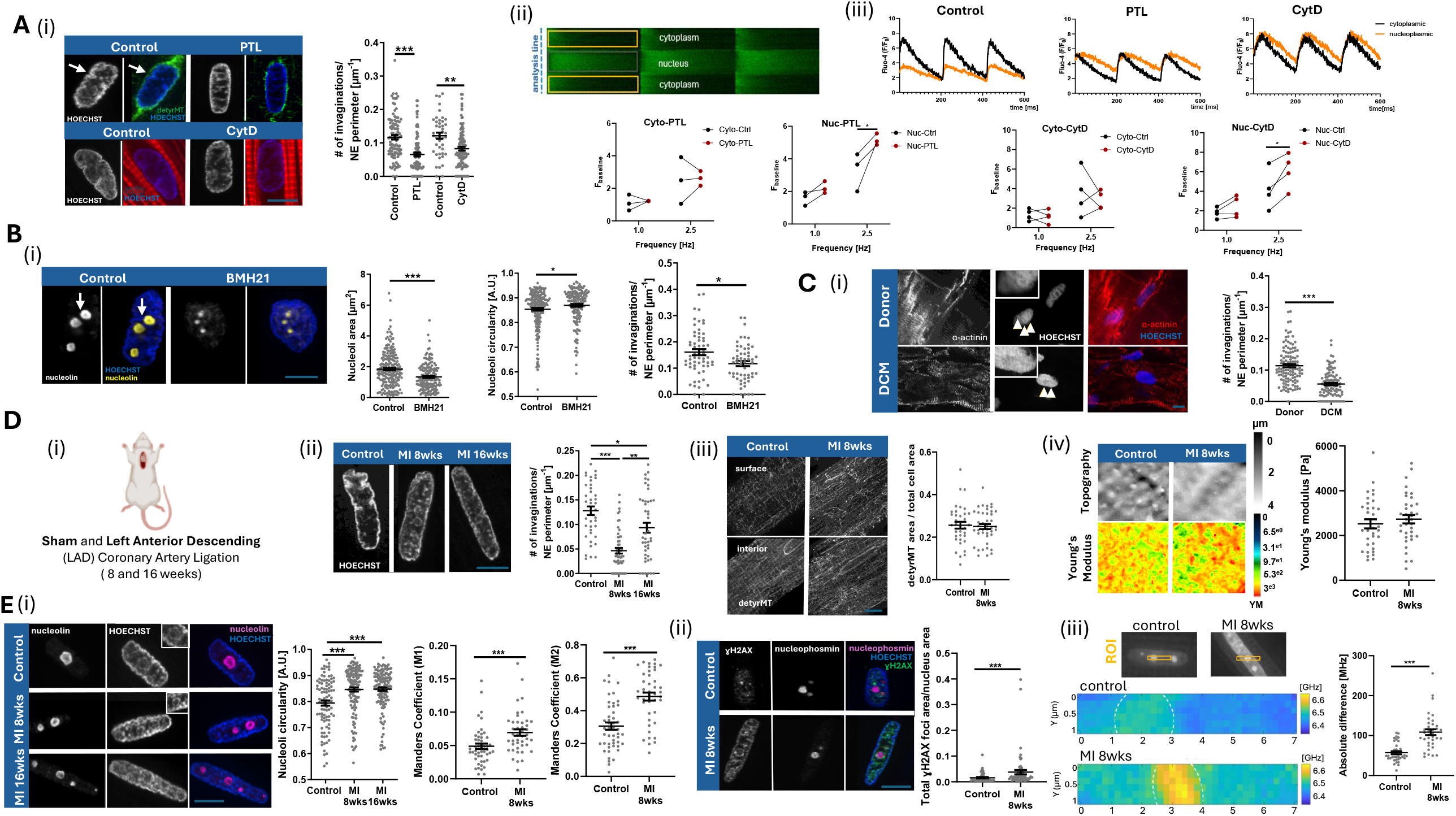
**Ai**, Left, representative immunocytochemistry micrographs of detyrMT (green), actin (red) and nuclei (HOECHST, blue). White arrow indicates detyrMT inside the NI. Right, scatter plot of NI quantification in control, PTL and CytD-treated cardiomyocytes (N=3-5 rats, 44-106 nuclei). **Aii**, Line-scan schematic of cytoplasmic and nucleoplasmic CaTs in a cardiomyocyte. **Aiii**, Representative recordings of distinct subcellular regions, as indicated in (ii), baseline quantification in control, PTL and CytD at 1 and 2.5Hz (bottom) (N=3-4 rats, 32-36 cells). **Bi**, Left, representative immunocytochemistry micrographs of nucleolin (yellow) and nuclei (HOECHST, blue). White arrow indicates NI-nucleoli localization. Right, scatter plots of area, circularity (N=5 rats,162-209 nucleoli) and NI number (N=3 rats, 60-57 nuclei) in control and BMH21-treated cells. **Ci**, Left, representative immunohistochemistry micrographs α-actinin (red) and nuclei (HOECHST, blue). Right, scatter plot of NI quantification in donor and DCM tissue (N=8 patients, 99-120 cells). **Di**, Animal model schematic. Representative immunocytochemistry micrographs of nuclei (HOECHST, grey) in **Dii** and detyrMT (grey) in **Diii**. Scatter plots of NI quantification (**Dii**) in control, 8-weeks and 16-weeks MI cardiomyocytes (N=3-4 rats, 40-44 cells) and detyrMT area quantification in control and 8-weeks MI cardiomyocytes (N=3 rats, 35-42 cells). **Div**, Left, mechano-Scanning Ion Conductance Microscopy representative topography and Young’s modulus maps. Right, Young’s modulus quantification in control and 8-weeks MI (N=3 rats, 33-36 cells). Representative immunocytochemistry micrographs of nucleolin (magenta) and nuclei (HOECHST, blue) in **Ei** and nucleophosmin (magenta), γH2AX-DNA damage (green) and nuclei (HOECHST, blue) in **Eii**. Scatter plots of circularity (N=3-4 rats, 96-137 nucleoli) and chromocenter Manders Coefficient (M1,M2) (N=3 rats, 43-49 cells) in control and MI cardiomyocytes (**Ei**) and γH2AX foci area (**Eii**) (N=3 rats, 48-63 cells). **Eiii**, Left, representative heatmap. Right, Brillouin shift absolute difference in control and 8-weeks cardiomyocytes (N=3 rats, 36-37 cells). Statistics: nested t-test (**Bi**,**Diii-iv**), unpaired t-test Kolmogorov-Smirnov (**Bi**,**Ci**,**Ei-iii**), non-parametric Kruskal Wallis with Dunn’s multiple comparison (**Ai**,**Dii**,**Ei**).Normality tested with Shapiro-Wilk.Data represent mean±SEM. Scale bar 5µm.

Consistent with what has been reported in other cell types, we found that some of the NIs, extend all the way to the nucleolus (Fig.Bi) (4). It has been shown that nuclear condensates, such as nucleoli, generate interfacial tension to produce piconewton forces capable of repositioning endogenous DNA loci and deforming the surrounding nuclear environment (1). To test a potential nucleolar contribution in NIs formation, we induced nucleolar stress with the RNA polymerase I inhibitor, BMH21. This reduced nucleolar area, increased nucleolar circularity and produced a decline in NIs. Thus, NI formation arises from coordinated forces generated both outside and inside the nucleus, with nucleoli acting as underappreciated mechanical drivers (Fig.Bi). This dual mechanism represents a conceptual advance in cardiomyocyte nuclear mechanics.

To determine whether NIs remodelling is relevant in disease, we examined human end-stage HF tissue. Consistent with earlier report, Dilated Cardiomyopathy (DCM) samples showed a depletion of NIs, indicating that NIs loss is a robust feature of terminal myocardial failure (Fig.Ci) (2).

Because nuclear abnormalities often arise early in disease, we asked whether NIs reduction also occurs at earlier stages. We utilized an 8- and 16-week Myocardial Infarction (MI) rat model (Fig.Di). NIs loss was already apparent at 8-weeks post-MI, well before end-stage remodelling. At this stage, detyrMT levels were unchanged and high-resolution, non-contact mechano-Scanning Ion Conductance Microscopy revealed only a non-significant increase in cardiomyocyte transverse stiffness, indicating that NIs loss cannot be attributed to detyrMT remodelling, which typically drives stiffness changes later in HF progression (Fig.Di-iv) (3). Instead, we observed nucleoli became more circular at 8 weeks post-MI, reflecting potentially altered interfacial tension at the nucleoli–NI border, in line with our nucleolar perturbation experiments. These nucleolar changes coincided with the emergence of prominent chromocentres (Fig.Ei zoom-in) surrounding nucleoli (M1 score for chromocenter-nucleolin overlap; M2 score for nucleolin-chromocentre overlap), indicating local reorganization of repressive chromatin, which overlapped with γH2AX DNA damage particularly in the nucleolar region (Fig.Ei-ii). Given that nucleoli can tether heterochromatin and impose structural constraints, chromocentre accumulation supports a shift in nucleolar–chromatin interactions during 8-week post-surgery HF.

These observations of nucleolar remodelling associated with NI loss may indicate a different nucleolar mechanical state. To test this, we analysed nucleolar biomechanics *in situ* using non-destructive, contact-free Brillouin Microscopy. Here, we measured the Brillouin shift (associated with changes in the longitudinal elastic modulus), showing statistically relevant increase in the relative values at 8-weeks post-MI, previously correlated with nucleoli stiffening in a target-specific manner (Fig.Eiii) (5). This altered mechanical contrast shows that nucleoli remodel before major cytoskeletal changes.

The first more detailed description of the double-membrane nuclear folds was published in 1997 by Fricker et al (6). However, the nuclear microstructure has not received much attention since that time. Only in the past decade, the NI found in different cell types, have drawn significant interest due to their potential role in nuclear architecture, gene regulation and cellular signalling processes (7).

Using state-of-the-art approaches, our study demonstrates that NI are regulated by the actin cytoskeleton, detyrosinated microtubules and nucleoli that tether to NI within the nucleus. We observe a decline in NI at 8- and 16-weeks post-MI. Importantly, we show that nucleolar remodelling precedes cytoskeletal abnormalities and drives the loss of NI at 8 weeks post-MI, repositioning nuclear and nucleolar integrity as central determinants of cardiac functional decline. Our findings identify the cardiomyocyte nucleolus as a previously underappreciated architect of nuclear organization and highlight nucleolar preservation as a promising therapeutic strategy to mitigate adverse cardiac remodelling.

## Materials and Methods

### Animal models

All animal surgical procedures and perioperative management were carried out in accordance with the United Kingdom Home Office Guide on the Operation of the Animals (Scientific Procedures) Act 1986 and EU Directive 2010/83, under the approval of the Animal Welfare and Ethics Review Board of Imperial College London. Myocardial infarction by proximal coronary ligation was used as a model for HF in adult rats as previously described (8). In brief, coronary ligation was performed on randomly selected male Sprague-Dawley rats to generate transmural infarcts and induce chronic HF. Rat models of 8- and 16-weeks post coronary artery ligation were used to represent gradual change in the heart performance leading to chronic HF. 16-weeks animals exhibit ventricular dilatation, reduced ejection fraction, as well as several structural, molecular, and electrophysiological changes. Both models were characterized here (8, 9).

### Cell isolation

Left ventricle cardiomyocytes were obtained from adult male Sprague-Dawley rats (250-300g). Cardiomyocyte isolation was done as previously described (10). Rats were anesthetized with 5% isoflurane and then killed by cervical dislocation. Hearts were extracted and placed in Tyrode solution containing: 140 mM NaCl, 6 mM KCl, 1mM MgCl2, 1mM CaCl2, 10mM glucose and 10mM HEPES, adjusted to pH 7.4 with 2 mmol/L NaOH. Hearts were placed in a Langendorff system after aortic cannulation, and perfused with Tyrode solution for 5 min, then with low Ca2+ oxygenated solution for 5 min: 120mM NaCl, 5.4mM KCl, 5mM MgSO4, 5mM sodium pyruvate, 20mM glucose, 20mM taurine, 10mM HEPES, 5mM nitrilotriacetic acid, and 0.04mM CaCl2, adjusted to pH 6.96 with 2 mmol/L NaOH, and finally for 10 min with enzyme solution: 120mM NaCl, 5.4mM KCl, 5mM MgSO4, 5mM sodium pyruvate, 20mM glucose, 20mM taurine, 10mM HEPES, and 0.2mM CaCl2, pH 7.4 with collagenase type 2 (1 mg/ml; Worthington) and hyaluronidase (0.6 mg/ml; Sigma-Aldrich). Cardiomyocytes were plated on dishes coated with laminin and allowed to attach to the dish/coverslip for at least 45 min before experiments. Cardiomyocytes were used on the same day of isolation. Cells isolated from the 8- and 16-week post-coronary artery ligation model were considered “failing” cells. These failing cells are isolated from the left ventricular wall of the infarcted heart after the infarcted area is removed.

### Cell treatment

The following drugs were used for the treatment of cardiomyocytes at 37°C: parthenolide (PTL) 10µM, cytochalasin D (CytD) 10µM and BMH21 1µM.

### Confocal Ca^2+^ imaging of nucleoplasmic and cytoplasmic CaTs

Cells were loaded with Fluo-4/AM (8 µmol/L; Thermo Fisher Scientific, USA). Simultaneous imaging of nucleoplasmic and cytoplasmic CaTs was performed in Fluo- 4/AM-loaded cardiomyocytes (excitation and emission wavelengths were 488 nm and >515 nm, respectively) using an inverted microscope equipped with a Plan Neofluar 40x/1.3 N.A. oil-immersion objective and a Zeiss LSM 510 Meta (Zeiss, Germany) confocal microscope. The pinhole was 1 Airy unit, resulting in an optical slice thickness of 0.9 μm. The confocal plane through the middle (z-axis) of the nuclei ensured nucleoplasm-exclusive signal collection. A 512-pixel line including the nucleus was scanned every 1.27 ms. Consecutive scan lines were stacked over time and analyzed using a custom-made MATLAB script (The MathWorks, Inc., USA). For quantification of subcellular CaTs cells were field-stimulated via platinum electrodes at 0.5 Hz. To measure the frequency-dependent changes, pacing rate was gradually increased from 1 Hz to 2.5 Hz.

### Immunocytochemistry

The cells were fixed in 4% paraformaldehyde for 10 minutes, permeabilized in Triton X-100 for 5 minutes and blocked for 0.5 hour in 5% Bovine Serum Albumin (BSA) as a blocking buffer. Next, the cells or tissues were incubated with a primary antibody overnight, washed with PBS and incubated for another hour with a secondary antibody and HOECHST. Primary antibodies used: anti-mouse α-actinin (Sigma, MA1-22863), anti-rabbit detyrosinated tubulin (Invitrogen, MA5-44591), anti-rabbit nucleolin (Cell Signaling, #14574), anti-mouse nucleophosmin (Abcam, ab10530), γH2AX (Cell Signaling, #5438). Secondary antibodies Alexa Fluor were all diluted 1:500. Wheat Germ Agglutinin (WGA; ThermoFisher, W32465) was used to stain the membrane, actin phalloidin (Biotium, 00044) and Hoechst 33342 (Thermo Scientific, 62249) was used to stain nuclei. Coverslips were mounted using Prolong Gold (ThermoFisher, P36930). Samples were imaged using Nikon Ti2 SoRa Spinning Disc Microscope with 60x TIRF objective, using a 1x or 2.8x SoRa disc. A Z-stack of images was taken from the top to the bottom of the cell in 1 μm steps.

### mechanoSICM

To map YM, cells were scanned using a modification of SICM in “hopping” mode (mechanoSICM) (11). Aerostatic, constant pressure (15 kPa) was delivered to the nanopipette (250-to 350-nm diameter) and propelled the inner solution, forming a hydrojet, applied throughout the scanning procedure (10 µm × 10 µm). The current– distance dependence was used to map the topography (0.7% of current reduction) and YM maps with two additional channels, SP1 and SP2, 1% and 2% current reduction, respectively. Topographical slope correction was introduced to correct for the uneven membrane surface. Between three and five measurements in each domain (z-groove and crest) were averaged per cell.

### Brillouin micro-spectroscopy

Brillouin micro-spectroscopy was used to assess the mechanical properties of nucleoli in fixed cells. A commercial system (LightMachinery) was integrated onto an inverted microscope (Nikon Eclipse Ti2-U) and operated with a 660 nm continuous-wave laser (Cobolt Flamenco, ∼100 mW). The beam passed through a temperature-stabilised air-spaced etalon and 0.3 OD attenuation and was coupled into a confocal circulator (LightMachinery HF-13311) with a polarising beam splitter and quarter-wave plate to produce circular polarisation. Excitation and signal collection were performed using a 40× water-immersion objective (Nikon Plan Apochromat LWD Lambda S 40×C WI).

Backscattered light was routed into a HyperFine spectrometer (LightMachinery HF-8999-PK-660) equipped with a “pump-killer” etalon providing ∼60 dB Rayleigh suppression. Brillouin Stokes and anti-Stokes components were dispersed by a VIPA and reflective echelle grating and detected simultaneously on a CMOS camera (Hamamatsu ORCA-Fusion C14440-20UP). A typical acquisition time was 250 ms with <1 pm spectral resolution.

Brillouin spectra were fitted using a damped harmonic oscillator (DHO) model to extract the Brillouin frequency shift (*v*_*B*_) and linewidth (Γ_*B*_):

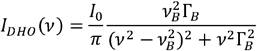

Scans were acquired with 0.2 µm step size. For each cell, the Brillouin shift was averaged over five pixels within the nucleolus and compared with five pixels in the surrounding nucleoplasm. Pixel positions were validated by overlaying Brillouin maps with DAPI (nucleus) and nucleolar immunofluorescence. A custom Python script was used for spectral fitting and regional averaging.

### Cryosectioning

Approval to work with donor and failing samples was obtained from NHS BT (REC reference:16/LO/1568) and Brompton Harefield& NHLI Ethics Committee under the Biobank REC (reference:09/H0504/104+5). Around 5-mm left ventricle Donor and Dilated Cardiomyopathy heart tissues were embedded in OCT. 10 µm thick cryosections were obtained on the cryostat microtome (Leica Biosystems) and further labelled with α-actinin and HOECHST.

### Quantification and statistical analysis

Analysis was made using ImageJ 1.54 software. Nuclear invaginations were quantified as the number of invaginations normalized to the nuclear perimeter, showing similar changes in control versus HF as here (2). All statistical analysis and graphs were performed/generated using GraphPad Prism 8. The Shapiro-Wilk test was used to test for dataset normality. Nested analyses were performed to assess significance; if data failed the normality test, non-parametric tests were applied. Data were represented as mean±SEM, *p<0.05, **p0<0.01, ***p<0.001

## Acknowledgments

The authors would like to thank Giedrė Astrauskaitė for Brillouin data analysis support and acknowledge the Cellular Mechanosensing and Functional Microscopy Centre at Imperial College London for access to equipment. Fig.D(i) was created with Biorender.com.

## Author contribution

Conceptualization, H.W., P.S., T.I., S.L.H.; Data curation H.W., I.M., P.S., A.N.K, J.F.; Investigation P.S., I.M., H.W., S.L.H.; Methodology I.M., T.I., S.L.H.; Writing-original draft H.W., P.S.; Writing-review&editing P.S., T.I., M.V., S.L.H., J.G.; Supervision P.S., T.I., M.V., S.L.H., J.G.

## Data availability

The data sets used and analyzed during the current study are available from the corresponding author on reasonable request.

